# Verification of human nucleotide sequence reagents and cell line identities in original circRNA articles published in high impact factor journals

**DOI:** 10.64898/2026.05.28.728608

**Authors:** Pranujan Pathmendra, Francisco J. Enguita, Jennifer A. Byrne

**Affiliations:** School of Medical Sciences, Faculty of Medicine and Health, The University of Sydney, NSW, Australia; Instituto de Genética, Faculdade de Medicina, Universidade de Lisboa, 1649-028 Lisboa, Portugal; NSW Health Statewide Biobank, NSW Health Pathology, Camperdown, NSW, Australia

**Keywords:** circRNA, cell lines, errors, nucleotide sequence reagents, non-verifiable

## Abstract

Numbers of research articles studying circRNAs have increased rapidly since 2017. Previous analyses of human circRNA articles in two high impact factor cancer research journals identified papers with wrongly identified nucleotide sequence reagents and circRNAs whose identities could not be independently verified. In the present study, verification of human nucleotide sequence reagent and cell line identities in retracted circRNA articles published from 2017-2021 in high impact factor journals found wrongly identified nucleotide sequences and/or cell lines in all 13 retracted papers. Similar analyses of human circRNA papers published in high impact factor journals in 2022 found wrongly identified, non-verifiable and/or questionable reagents in 71% (84/118) papers, where 51% (60/118) papers described at least one wrongly identified reagent. When individual error types and features of concern were considered, 2022 circRNA papers described wrongly identified nucleotide sequence reagents (52/118, 44%), questionable circRNA probes that did not meet accepted targeting requirements (34/118, 29%), non-verifiable nucleotide sequences (25/118, 21%), wrongly identified cell lines (22/118, 19%), and/or non-verifiable cell line identifiers (6/118, 5%). In summary, wrongly identified, non-verifiable and/or questionable reagents were unexpectedly frequent in human circRNA papers in high impact journals, highlighting the need for critical engagement with the circRNA literature.

## Introduction

The analysis of gene functions relies on laboratory experiments in model systems. With over 40,000 human genes and many hundreds of thousands of associated linear and circular transcripts, the task of understanding human gene functions requires both scaled and targeted experimental approaches. Gene function experiments can be expensive, technically demanding and slow (Byrne et al. 2022), and challenging to reproduce (Yan et al. 2019; Errington et al. 2021). The combination of technical and structural barriers to the reproduction of gene function experiments means that the results of many gene function studies are likely to remain unchallenged within the literature, despite potentially being implausible, non-generalisable, or simply incorrect.

Several research groups have highlighted publications in non-coding RNA (ncRNA) biology that may describe unreliable results. Some papers have analysed ncRNAs in relation to proposed targets where the predicted stoichiometries of regulators and targets do not predict biologically relevant functions (Kilikevicius et al. 2022). Human microRNAs have been studied in cell lines that are not indicated to express targeted microRNAs at detectable levels (Kent et al. 2014; Kachooei et al. 2025), and the status of many claimed microRNAs has been questioned (Hackenburg et al. 2025; Orang et al. 2025).

The validity of results from gene function experiments also relies on reagents that correctly target claimed genes or transcripts of interest, and cell line models that represent claimed cell types or diseases (Byrne, 2025). Conversely, implausible or incorrect experimental results can become apparent by verifying experimental reagent identities. The lack of visual sense in both short nucleotide sequences and cell line identifiers can mask experimental reagents that have been wrongly identified, leading to data associated with wrongly identified reagents persisting in the literature for decades (Oste et al. 2024). Published examples of wrongly identified nucleotide sequence reagents have been recognised for at least 26 years (Jung et al. 2000), whereas wrongly identified human cell line models have been recognised for almost 60 years, initially through the discovery of cell line cross-contamination by HeLa cells (Korch and Varella-Garcia, 2018).

Our team has described human gene research papers with wrongly identified and non-verifiable reagents across a range of journals (Byrne, 2025). We initially found incorrect non-targeting sequences in papers that described the effects of knocking down human single genes in cancer cell lines (Byrne and Labbé, 2017; Labbé et al. 2019). We then studied papers that described the functions of human ncRNAs (Park et al. 2022; Oste et al. 2024) and analysed gene functions in the context of cancer chemotherapy sensitivity or resistance (Park et al. 2022). Most papers with wrongly identified nucleotide sequence reagents and cell lines were published in low to moderate impact fact (IF) journals (Byrne and Labbé, 2017; Labbé et al. 2019; Park et al. 2022; Oste et al. 2024), and screening original papers in the journals *Gene* and *Oncology Reports* found proportions of papers with wrongly identified nucleotide sequences ranging from 0.5-4.1% (*Gen*e, 2007-2018) to 8.3-12.6% (*Oncology Reports*, 2014-2018) (Park et al. 2022). Analyses of 2021-2023 papers studying human *miR-145* found articles that described both wrongly identified nucleotide sequences and cell lines, as well as claimed human cell lines that may not exist (Oste et al. 2024).

The unexpected frequency of wrongly identified reagents in journals of low to moderate IF could reflect the peer review and publishing standards of these journals and/or that most scientific journals are of low to moderate IF (Siler and Larrivière, 2022). We therefore verified nucleotide sequence reagent identities in papers in two high IF cancer journals, *Molecular Cancer* and *Oncogene*, defined as journals with a 2019 IF≤ 7 (Pathmendra et al. 2024). Screening all original *Molecular Cancer* papers in 2014, 2016, 2018, and 2020 revealed rising proportions of papers with wrongly identified nucleotide sequences, up to 38% original *Molecular Cancer* papers in 2020 (Pathmendra et al. 2024). As some 2018 and 2020 *Molecular Cancer* papers analysed human circRNAs, we verified nucleotide sequence identities in 2020 *Oncogene* papers that studied human miRNAs and/or circRNAs, where wrongly identified nucleotide sequences were found in 40% screened papers (Pathmendra et al. 2024). We also found circRNA targeting reagents whose identities could not be verified, as the claimed human circRNA could not be independently identified using publicly available data (Pathmendra et al. 2024). This finding echoed earlier descriptions of non-verifiable (NV) human circRNAs in cancer research publications by Patop and Kadener (2018).

As our research had identified human circRNA publications that described wrongly identified nucleotide sequence reagents and/or cell lines, as well as NV circRNAs or human cell line identifiers (Pathmedra et al. 2024; Oste et al. 2024), the present study verified reagent identities in new corpora of human circRNA articles. We verified human nucleotide sequence and cell line identities in (i) retracted human circRNA papers published in high IF journals between 2017-2021, where high IF was defined as 2022 IF≥7, and (ii) human circRNA papers published in high IF journals in 2022. As we will describe, we found that all 13 retracted circRNA papers in high IF journals described wrongly identified nucleotide sequence reagents and/or cell lines, where only one retraction notice recognised any wrongly identified reagents. Similar analyses of 2022 circRNA papers in high IF journals found wrongly identified, non-verifiable and/or questionable reagents in 71% (84/118) papers, where 51% (60/118) papers described at least one wrongly identified nucleotide sequence reagent and/or cell line. We will discuss the significance of these results in terms of the reliability of human circRNA research published in high IF journals.

## Results

### Wrongly identified reagents in retracted circRNA papers in high impact factor journals

We found 93 retracted papers in 32 journals that referred to human circRNAs in their titles (see Methods). Retracted papers were published between August 2017-January 2021 and retracted between December 2018-July 2023. Thirteen (14%) retracted circRNA papers were published in 6 high IF journals (2022 IF≥7) and were analysed further. We will refer to these 13 retracted circRNA papers as the retracted circRNA corpus.

All 13 retracted papers were authored by teams from China, where most were affiliated with hospitals (Table 1). Almost all (12/13) retracted papers studied a single cancer type, with the remaining paper (PMID 31900142) studying circRNA in the context of atherosclerosis. Several retracted circRNA papers included the terms “axis” (n=5) or “sponge” (n=3) in their titles (Table S1). Retraction times for the 13 papers ranged from 6-55 months after publication (Table S1).

**Table 1.** Summary of retracted circRNA and 2022 circRNA publication corpora.

|  | **Retracted circRNA** | **2022 circRNA** |
| --- | --- | --- |
| Papers n= | 13 | 118 |
| Publication year(s) | 2017-2021 | 2022 |
| Journals n= | 6 | 47 |
| Publishers n= | 2 | 8 |
| Journal IFs, median (range) | 9.7 (range) | 9 (range) |
| Proportion (%) papers authored by teams from China | 13/13 (100%) | 98/118 (83%) |
| Proportion (%) papers affiliated with hospitals | 12/13 (92%) | 80/118 (68%) |
| Proportion (%) papers with nucleotide sequence reagents | 12/13 (92%) | 113/118 (96%) |
| Nucleotide sequence reagents analysed n= | 254 | 3,370 |
| Nucleotide sequence reagents analysed per paper, median (range) | 23 (2-55) | 22 (1-154) |
| Proportion (%) wrongly identified nucleotide sequences | 42/254 (17%) | 197/3,370 (5.8%) |
| Proportion (%) papers with wrongly identified nucleotide sequences | 11/13 (85%) | 52/113 (46%) |
| Wrongly identified nucleotide sequences per paper, median (range) | 1 (1-4) | 2 (1-32) |
| Proportion (percentage) papers with human cell lines | 13/13 (100%) | 90/118 (76%) |
| Human cell line descriptions analysed n= | 52 | 392 |
| Human cell lines analysed per paper, median (range) | 4 (1-8) | 4 (1-10) |
| Proportion (%) human cell line descriptions | 6/52 (12%) | 57/392 (15%) |
| Wrongly identified human cell lines n= | 6 | 27 |
| Proportion (%) papers with wrongly identified cell lines | 3/13 (23%) | 22/90 (24%) |
| Wrongly identified cell lines per paper, median (range) | 1 (1-4) | 3 (1-7) |
| Proportion (%) problematic papers (wrongly identified nucleotide sequences and/or human cell lines) | 13/13 (100%) | 60/118 (51%) |
| Proportion (%) problematic papers authored by teams from China | 13/13 (100%) | 54/60 (90%) |
| Proportion (%) problematic papers from China affiliated with hospitals | 12/13 (92%) | 43/54 (80%) |

We verified the identities of 254 nucleotide sequence reagents across 12 retracted circRNA papers, which involved checking a median of 23 nucleotide sequences per paper (range: 2-55, n=12 papers) (Table 1). Seventeen percent (42/254) nucleotide sequence reagents were predicted to be wrongly identified (Table 1), which were distributed across 11 retracted papers, with a median of 1 incorrect nucleotide sequence per paper (range: 1-4, n=11 papers) (Table 1). Most (33/42, 79%) wrongly identified nucleotide sequences were predicted to target a different human gene or genomic sequence from that claimed. Many wrongly identified reagents (29/42, 69%) were claimed to target circRNAs (Table 2). Two retracted circRNA papers with wrongly identified nucleotide sequences described a total of 14 non-verifiable (NV) circRNA targeting reagents, where the claimed circRNAs could not be identified (see Methods) (Table 3, Table S1).

**Table 2.** Wrongly identified nucleotide sequence reagents in retracted circRNA and 2022 circRNA corpora.

|  | **Retracted circRNA** | **2022 circRNA** |
| --- | --- | --- |
| Proportion (%) wrongly identified nucleotide sequence reagents | 42/254 (17%) | 197/3,370 (5.8%) |
| Proportion (%) wrongly identified nucleotide sequences according to reagent type |  |  |
| RT-PCR primers | 36/42 (86%) | 130/197 (66%) |
| Single probes | 6/42 (14%) | 67/197 (34%) |
| Proportion (%) wrongly identified reagents according to error types |  |  |
| Claimed targeting reagents predicted to target other human gene/ genomic sequences | 33/42 (79%) | 116/197 (59%) |
| Claimed targeting reagents predicted to be non-targeting in human | 9/42 (21%) | 78/197 (40%) |
| Claimed non-targeting reagents predicted to target a human transcript | 0/42 (0%) | 3/197 (1.5%) |
| Proportion (%) wrongly identified reagents according to claimed target |  |  |
| Protein-coding genes (including gene promoters) | 5/42 (12%) | 88/197 (45%) |
| ncRNAs | 8/42 (19%) | 39/197 (20%) |
| circRNAs | 29/42 (69%) | 67/197 (34%) |
| Claimed circRNA targeting reagents predicted to target other human gene/ transcript/ genomic sequence | 25/29 (86%) | 50/67 (75%) |
| Predicted to not discriminate between circular and linear transcripts from claimed host gene | 3/25 (12%) | 36/50 (72%) |
| Predicted to target sequence shared by circRNA and linear transcript | 1/3 (33%) | 15/36 (42%) |
| Claimed divergent RT-PCR primers predicted to function as convergent primers | 0/3 (0%) | 8/36 (22%) |
| Predicted to map to BSJ and also target linear transcript | 2/3 (67%) | 13/36 (36%) |
| Predicted to target other gene/ transcript/ genomic sequence | 22/25 (88%) | 13/50 (26%) |
| Predicted to target different circRNA from that claimed | 2/22 (9%) | 8/13 (62%) |
| Predicted to target a different circRNA isoform from host gene | 0/22 (0%) | 2/8 (25%) |
| Predicted to target other gene/ transcript (not circRNA) | 20/22 (91%) | 5/13 (38%) |
| Duplicated RT-PCR primer (same sequence for forward and reverse) | 0/25 (0%) | 1/50 (2%) |
| Claimed circRNA-targeting reagents predicted to be non-targeting in human | 4/29 (14%) | 17/67 (25%) |
| Claimed circRNA targeting reagents within insufficient matches across BSJ | 1/4 (25%) | 3/17 (18%) |

**Table 3.** Non-verifiable (NV) nucleotide sequence reagents in retracted circRNA and 2022 circRNA corpora.

|  | **Retracted circRNA** | **2022 circRNA** |
| --- | --- | --- |
| Proportion (%) NV sequence reagents | 14/254 (5.5%) | 88/3,370 (2.6%) |
| Proportion (%) papers with verified nucleotide sequence reagents that described NV sequences | 2/12 (17%) | 25/113 (22%) |
| Proportion (%) papers that also described wrongly identified nucleotide sequences | 0/2 (0%) | 10/25 (40%) |
| Proportion (%) NV reagents according to reagent type |  |  |
| RT-PCR primers | 2/14 (14%) | 66/88 (75%) |
| Single probes (sgRNA, shRNA, siRNA, other oligonucleotide probes) | 12/14 (86%) | 22/88 (25%) |
| NV sequences per paper, median (range) | 7 (1-13) | 2 (1-14) |
| Proportion (%) NV reagents claimed to target circRNAs | 14/14 (100%) | 86/88 (98%) |
| Claimed circRNA identity could not be verified on circATLAS/ circBASE (authors did not provide or reference complete circRNA sequence) | 1/14 (7.1%) | 41/86 (48%) |
| No reference to specific circRNA transcript | 1/1 (100%) | 33/41 (80%) |
| Claimed circRNA target with unique identifier | 0/1 (0%) | 8/41 (20%) |
| Use of ArrayStar ID with no additional information, numeric component of circRNA ID did not correspond to any circBASE ID | 13/14 (93%) | 32/86 (37%) |
| Claimed circRNA ID resembled ArrayStar ID, no reference to ArrayStar in article methods | 0/14 | 11/86 (13%) |
| Unclear annotation of reagent sequence | - | 2/86 (2.3%) |
| Proportion (%) NV reagents not claimed to target circRNAs | 0/14 (0%) | 2/88 (2.3%) |
| Short nucleotide sequence (n=7 nucleotides) | - | 1/2 (50%) |
| Unclear annotation of reagent sequence | - | 1/2 (50%) |

We also verified the identities of 52 human cell line descriptions in the retracted circRNA corpus (Table 1). Retracted papers described a median of 4 cell lines per paper (range: 1-8, n=13 papers) (Table 1). Wrongly identified cell lines were found in 3 retracted papers, where one paper (PMID 30546088) also described a wrongly identified nucleotide sequence (Table S1). Almost all problematic cell lines (5/6) were contaminated, either by HeLa cells (BGC-823, MGC-803, SGC-7901) or another cell line of the same cancer type (K1, MKN-28).

### Analysis of retraction notices and associated PubPeer commentary

While all 13 retracted circRNA papers included wrongly identified nucleotide sequence reagent(s) or cell lines, only one retraction notice (PMID 37101243) mentioned wrongly identified reagents, recognising the use of four contaminated cell lines (BGC-823, MGC-803, MKN-28, SGC-7901) that this study also identified (Table S1). Instead, most (10/13) retraction notices cited image integrity concerns as reasons for retraction. The retraction notices for 2 circRNA papers published by *Journal of Experimental and Clinical Cancer Research* and *Molecular Therapy Nucleic Acids* in 2020 (PMID 36691006, PMID 35592506) mentioned undeclared third-party involvement as a key concern undermining confidence in study data.

We also analysed post-publication comments for the retracted circRNA corpus on PubPeer (Barbour and Stell, 2020). Nine retracted papers received PubPeer comments that were mentioned in subsequent retraction notices, with a median time of 7 months (range: 3-19, n=9) between the first PubPeer comment that was relevant to the retraction and the subsequent retraction (Table S1). There was no mention of wrongly identified nucleotide sequences or problematic cell lines in any PubPeer comments for retracted circRNA papers. In contrast, almost all (9/10) retraction notices that referred to image integrity concerns had received earlier PubPeer comments describing image integrity concerns (Table S1).

### Human circRNA articles in high IF journals

We identified 7,913 human circRNA papers published from 2001-2025 that were available via the Web of Science (Figure 1). Numbers of human circRNA papers remained below 100 papers/ year until 2017, after which the numbers of circRNA papers per year rose steeply, reaching over 1,200 papers in 2020-2022 (Figure 1).

**Figure 1.**
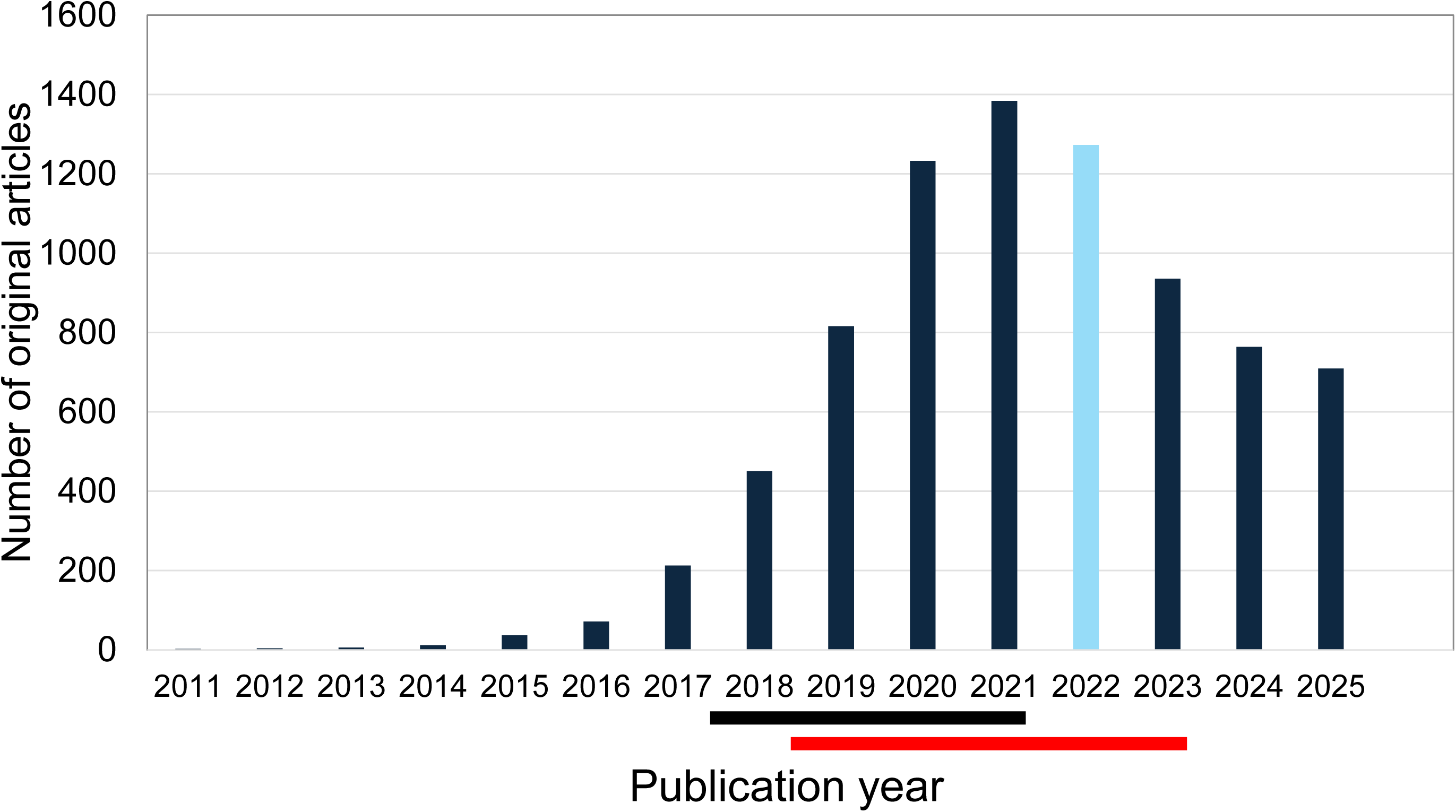
Number of original circRNA papers (Y axis) published from 2001 to 2022 according to publication year (X axis). Papers published in 2022 are highlighted in light blue. Horizontal bars below the X axis indicate when papers in the retracted circRNA corpus were published (black) or retracted (red), respectively.

We analysed circRNA papers published in 2022, as the most recent year available for analysis when this study commenced. In 2022, 1,273 original circRNA papers were published across 475 journals, of which 98% (464/475) were assigned a 2022 journal IF. Fifteen percent (190/1,276) original 2022 circRNA papers were published in journals with a 2022 IF≥7. Most (118/190, 62%) papers met the study inclusion criteria, in that they described original experiments, studied the function of at least one human circRNA, and either described the sequence of at least one nucleotide sequence reagent claimed to target a wild-type human transcript or genomic sequence, or at least one human cell line. We refer to these 118 human circRNA papers in high IF journals in 2022 as the 2022 circRNA corpus.

Papers in the 2022 circRNA corpus had been collectively cited 3,603 times according to Google Scholar by 25 September 2024. Most papers in the 2022 circRNA corpus (98/118, 83%) were authored by teams in China (Table S2). More papers from China were affiliated with hospitals (79/98, 81%), compared with papers from other countries (1/20, 5%) (Fisher’s Exact Test, p<0.0001, n=118).

### Wrongly identified nucleotide sequence reagents in the 2022 circRNA corpus

Almost all 2022 circRNA papers (113/118, 96%) described nucleotide sequence reagents whose identities could be verified (see Methods) (Table 1). These 113 papers described 3,370 nucleotide sequence reagents, with a median of 22 nucleotide sequences per paper (range: 1-154, n=113 papers) (Table 1). Overall, 197 (5.8%) nucleotide sequences were predicted to be wrongly identified (Table 1). Most were targeting reagents that were predicted to target a different human gene or genomic sequence from that claimed (116/197, 59%), followed by targeting reagents that were predicted to be non-targeting in human (78/197, 40%) (Table 2). Few reagents (3/197, 1.5%) were claimed non-targeting controls that were predicted to target a human transcript (Table 2).

Wrongly identified nucleotide sequences were distributed across 46% (52/113) 2022 circRNA papers, with a median of 2 wrongly identified nucleotide sequences/ paper (range: 1-32, n=52 papers) (Table 1). The numbers of nucleotide sequences screened and numbers of wrongly identified sequences per paper were weakly positively correlated (Spearman Rho=0.2807, 95% CI: 0.0002216 to 0.5203, p=0.0438, n=52).

### Wrongly identified circRNA targeting reagents in the 2022 circRNA corpus

One third of wrongly identified nucleotide sequences represented circRNA targeting reagents (67/197, 34%), which were found in 26/52 (50%) 2022 circRNA papers with wrongly identified nucleotide sequences (Table 2). Most (50/67, 75%) wrongly identified circRNA reagents were predicted to target a different human gene/genomic sequence from that claimed, where most reagents (36/50, 72%) were indicated to not discriminate between circular and linear transcripts (Table 2). This included reagent sequences that were predicted to be shared by circular and linear transcripts from the claimed host gene (15/36, 42%), sequences that mapped to the claimed BSJ but also matched a linear transcript (13/36, 36%), and claimed divergent RT-PCR primers that were predicted to function as convergent primers (8/36, 22%) (Table 2). Some incorrect circRNA reagents were predicted to be non-targeting in human (17/67, 25%), most of which were single probes (12/17, 75%) (Table 2, Table S3).

### Non-verifiable nucleotide sequence reagents in the 2022 circRNA corpus

Twenty-five 2022 circRNA papers (22%) were found to describe nucleotide sequences whose identities could not be verified (see Methods) (Table 3). We found 88 (88/3,370, 2.6%) NV nucleotide sequences, with a median of 2 NV nucleotide sequences per paper (range: 1-14, n=25 papers) (Table 3). Most NV nucleotide sequences were RT-PCR primers (66/88, 75%), with the remainder being single probes (Table 3).

Almost all (86/88, 98%) NV sequences were claimed to target human circRNAs (Table 3). Some NV sequences were linked with circRNA identifiers with insufficient information about the BSJ, such that we could not identify a specific transcript using circAtlas or circBase (41/86, 48%). Some NV circRNA identifiers appeared to be unique to the paper under study (8/41, 20%) (Table 3, Table S4), whereas other NV identifiers referred only to the host gene (33/41, 80%), for example *circGAPDH*. We also noted the use of ArrayStar circRNA identifiers (32/86, 37%) or identifiers that resembled ArrayStar identifiers (11/86, 13%) that could not be independently verified (Table 3, Table S4).

### circRNA targeting F/ISH and pull-down probes

We found 28 circRNA Fluorescence In Situ Hybridisation (FISH) or ISH probes in 27 papers (Figure 2A) and 30 RNA pull-down probes in 23 papers (Figure 2B) that could be mapped across the claimed BSJ. None of the 28 F/ISH probes met the targeting criteria of 22-24 nucleotide matches to each side of the BSJ (Zirkel and Papantonis, 2018) (Table S5), although one probe approximated these requirements (Figure 2A). Thirteen (43%) pull-down probes that mapped across the claimed BSJ satisfied the targeting criteria for circRNA pull-down reagents (Das et al. 2021; Gabryelska et al. 2024) (Table S5), where probes were 15-40 nt in length and showed equivalent matches to each side of the claimed BSJ (Figure 2B). Five circRNA pull-down probes exceeded 40 nt in length but showed similar matches (differing by 0-3 nt) to each side of the claimed BSJ (Figure 2B). Questionable F/ISH/ pull-down probes were described in 34/118 (29%) 2022 circRNA papers (Table S6).

**Figure 2.**
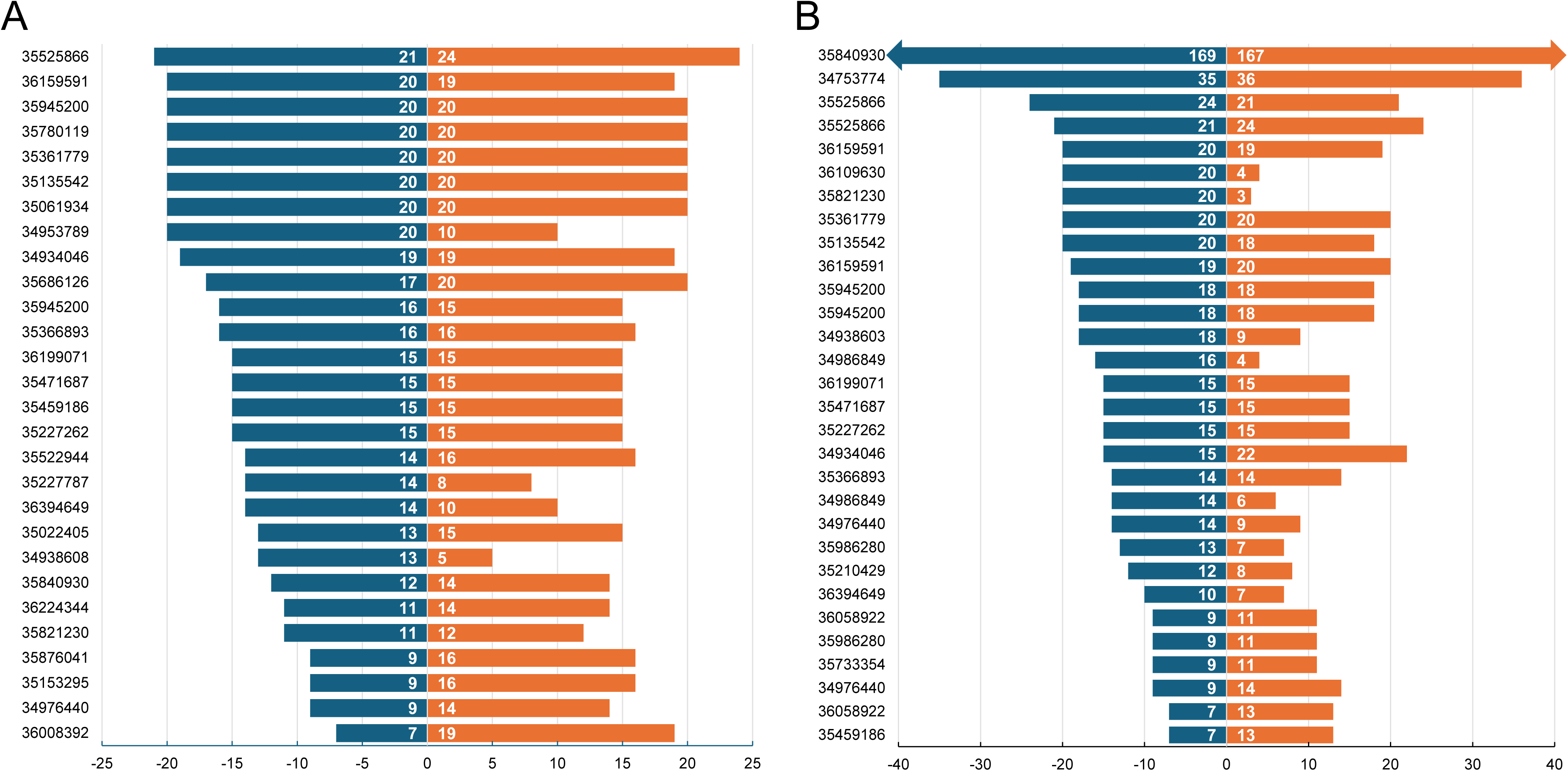
Human circRNA targeting (**A**) Fluorescence or in situ hybridisation (F/ISH) probes or (**B**) RNA pull-down probes described in 2022 circRNA papers that could be mapped to the claimed BSJ. (**A, B**) Probes are labelled according to the corresponding PMID (Y axis) and ranked according to lengths of 5’ sequence matches (negative numbers, X axis), where the position of the BSJ is shown as 0. Sequence matches for 5’/ 3’ probe sequences are represented by dark blue/ orange horizontal bars, respectively, with match lengths shown in white text. One pull-down probe (**B**) is partially shown, indicated by arrows.

### Wrongly identified and non-verifiable human cell lines in the 2022 circRNA corpus

We found 392 human cell line descriptions across 90 (76%) 2022 circRNA papers (Table 1). A median of 4 human cell lines were employed per paper (range: 1-10, n=90 papers) (Table 1). Fifteen percent (57/392) of human cell line descriptions corresponded to wrongly identified cell lines (Table 1), representing 27 wrongly identified cell lines distributed across 22 papers.

Most (44/57, 77%) descriptions of wrongly identified cell lines represented cross-contaminated cell lines, typically by HeLa cells as either the sole contaminant or with another unknown cell line (Table 4, Table S7). The most frequent HeLa-contaminated cell line was L02, wrongly claimed as a liver cell line (Table S7). Remaining descriptions referred to misclassified cell lines (13/57, 23%), such as HepG2 cells that were wrongly claimed to model hepatocellular carcinoma (Table 4, Table S7).

**Table 4.** Wrongly identified human cell lines in retracted circRNA and 2022 circRNA corpora.

|  | **Retracted circRNA** | **2022 circRNA** |
| --- | --- | --- |
| Proportion (%) papers that described human cell lines | 13/13 (100%) | 90/118 (76%) |
| Human cell line descriptions analysed n= | 52 | 392 |
| Proportion (%) cell line descriptions corresponding to wrongly identified cell lines | 6/52 (12%) | 57/392 (15%) |
| Proportion (%) papers with wrongly identified cell lines | 3/13 (23%) | 22/90 (24%) |
| Wrongly identified cell lines per paper, median (range) | 1 (1-4) | 3 (1-7) |
| Proportion (%) cross-contaminated human cell lines | 5/6 (83%) | 44/57 (77%) |
| HeLa cross-contaminated | 3/5 (60%) | 40/44 (91%) |
| HeLa as sole contaminant | 2/3 (67%) | 24/40 (60%) |
| HeLa and unknown cell line contamination | 0/0 (0%) | 13/40 (33%) |
| HeLa and cell line of claimed identity forming hybrid cell line | 1/3 (33%) | 3/40 (8%) |
| Cross-contamination by other cell line of claimed cancer type | 2/5 (40%) | 3/44 (7%) |
| Partial cross-contamination by non-human cells | 0/0 (0%) | 1/44 (2.3%) |
| Proportion (%) misclassified human cell lines | 1/6 (17%) | 13/57 (23%) |
| Hepatoblastoma claimed as hepatocellular carcinoma | 1/1 (100%) | 7/14 (50%) |
| Claimed hepatocellular carcinoma, likely derived from endothelial cells | 0/0 (0%) | 3/14 (21%) |
| Mismatch between RRID (research resource identifier) and claimed identity | 0/0 (0%) | 3/14 (21%) |
| Proportion (%) wrongly identified cell line descriptions according to claimed identity |  |  |
| Human cancer | 6/6 (100%) | 51/57 (89%) |
| Head and neck | 1/6 (17%) | 19/51 (37%) |
| Liver | 1/6 (17%) | 15/51 (29%) |
| Gastric | 4/6 (67%) | 11/51 (22%) |
| Lung | 0/0 (0%) | 3/51 (6%) |
| Oesophageal | 0/0 (0%) | 2/51 (3.9%) |
| Neuroblastoma | 0/0 (0%) | 1/51 (2.0%) |
| Untransformed cell lines | 0/0 (0%) | 6/57 (11%) |

We also found 6 human cell line identifiers (1.5%) whose identities could not be independently verified (see Methods) (Table 5). Five NV identifiers (CEC, HEB, HFC, MKN-87, SW-308) had not been previously described, whereas GSE-1 was recognized as an NV cell line by Oste et al. (2024). Similarly named cell lines were found for 4 NV identifiers (GSE-1, HFC, MKN-87, SW-408) (Table 5, Table S8). All 6 NV identifiers were found in at least 10 sources indexed by Google Scholar in May 2026 (Table 5, Table S8).

**Table 5.** Non-verifiable (NV) human cell line identifiers in the 2022 circRNA corpus ranked in alphabetical order.

| **NV cell line identifier<sup>a</sup>** | **2022 circRNA paper PMID/ DOI** | **Assumed/ claimed identity** | **Misspelled identifier according to Cellosaurus?** | **Similarly named cell line- identity**<br>Letters/ numbers in bold differ in NV identifier | **Google Scholar results**<br>n= (search string) May 2026 | **NV identifier indexed** by claimed external repository/ ATCC/ CellBank Australia?<sup>b</sup> (Y/N) |
| --- | --- | --- | --- | --- | --- | --- |
| CEC | https://doi.org/10.1016/j.snb.2022.132893<sup>c</sup> | Endothelial cells | No | Unclear | 38 ("CEC cell line") | N |
| **GSE1** | 35780119<sup>c</sup> | Gastric epithelial | No | **GES**-1- contaminated | 13 ("GSE1 cells") | N |
| HEB | 35927233<sup>c</sup> | Glial cell | No | Unclear | 85 ("HEB cell line") | N |
| HFC | 35101080<sup>c</sup> | Colonic epithelial | No | **FHC** - colonic epithelial | 52 ("HFC cell line") | N |
| MKN87 | 35354791<sup>c</sup> | Gastric cancer | No | Possible misspelling of **MNK-45** | 15 ("MKN87") | N |
| SW408 | 35184406 | Colorectal cancer | No | SW48 - colorectal cancer | 56 ("SW408") | N |

<sup>a</sup>NV identifier described by Oste et al. (2024) is shown in bold
<sup>b</sup>All repository catalogue searches used variations (+/- space/hyphen) of NV cell line identifiers, where relevant
<sup>c</sup>Papers also described wrongly identified reagent(s)

### Summary of reagent errors/ concerns for the 2022 circRNA corpus

Overall, 71% (84/118) 2022 circRNA papers included one or more incorrect, NV and/or questionable reagent, where 49% (41/84) papers described more than one error type or feature of concern (Figure 3, Table S9). While substantial overlap was noted between the 5 publication groups, almost all papers describing wrongly identified (20/22, 91%) or NV cell lines (5/6, 83%) included at least one error type or feature of concern (Figure 3).

**Figure 3.**
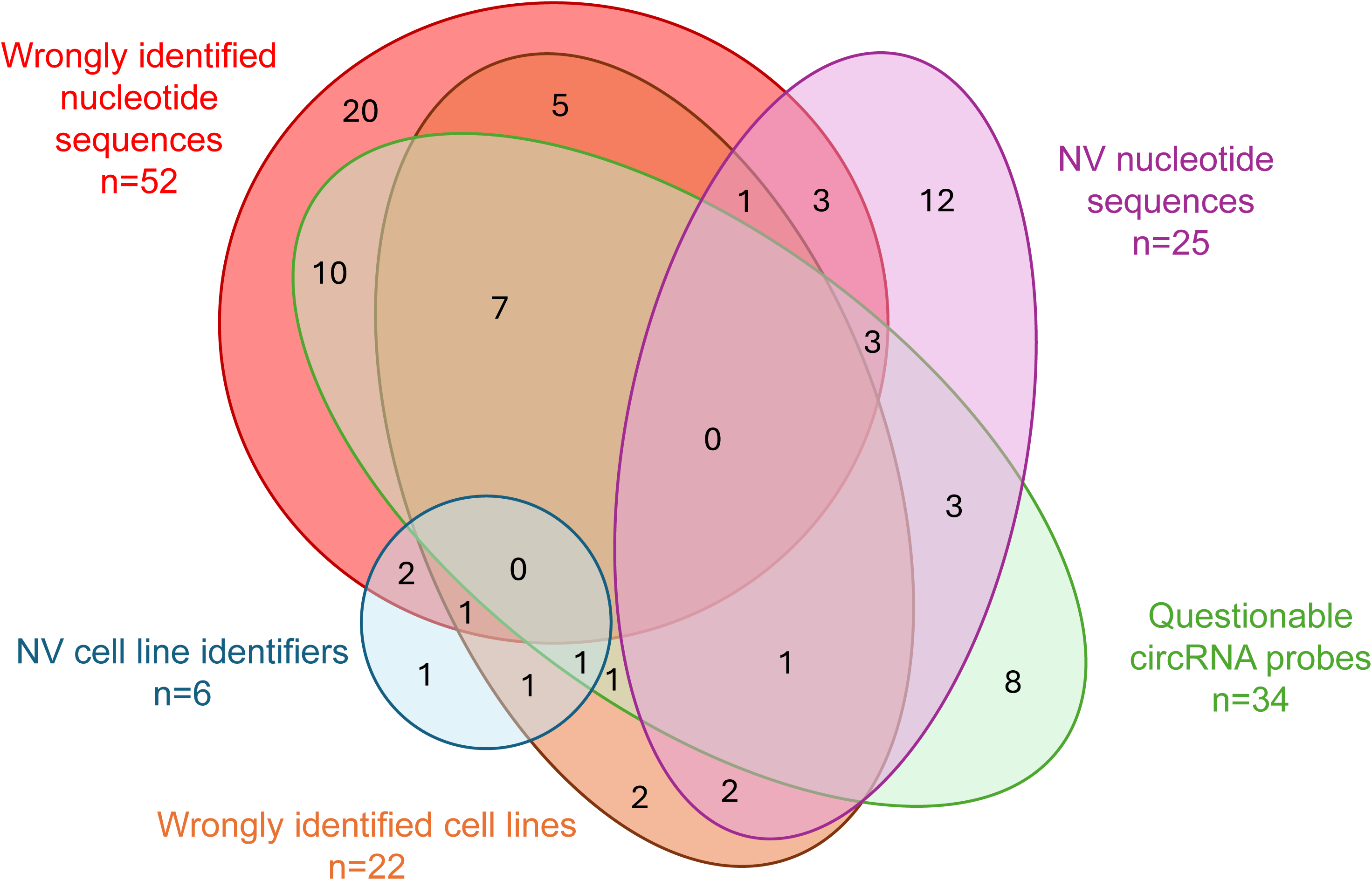
Venn diagram of 2022 circRNA papers in high IF journals with wrongly identified nucleotide sequences (red), wrongly identified cell lines (orange), NV cell line identifiers (blue), NV nucleotide sequences (purple) and/or questionable circRNA probes (green). NV = non-verifiable. Numbers show papers with corresponding features. Shape/ overlap sizes are for illustrative purposes only.

### Problematic 2022 circRNA papers

Half (60/118, 51%) 2022 circRNA papers described at least one wrongly identified reagent (nucleotide sequence and/or cell line) (Table 1, Figure 3). These papers are described as problematic, as they include verifiable errors. Some problematic papers claimed to study circRNAs in the context of a “gene axis” (14/60, 23%), or as “sponges” of miRNAs (10/60, 17%) (Table S2).

Problematic 2022 circRNA papers were published across 31 high IF journals with a median 2022 IF of 9.7 (range: 7-39.3, n=31 journals) (Table S2), where most (50/60, 83%) were published open access. Almost half (15/31, 48%) journals were published by Springer Nature, including *Cell Death and Disease* (2022 IF: 9), *Cell Death Discovery* (2022 IF: 7), *Journal of Translational Medicine* (2022 IF: 7.4), *Molecular*

*Cancer* (2022 IF: 37.3) and *Nature Communications* (2022 IF: 16.6) (Table S2). Problematic papers were largely authored by teams from China (54/60, 90%), where more problematic papers from China were affiliated with hospitals (43/54, 80%), compared with papers from other countries (1/6, 17%) (Fisher’s Exact Test, p=0.004, n=60 papers).

### Citations of problematic 2022 circRNA papers

The 60 problematic 2022 circRNA papers had been cited 1,559 times according to Google Scholar on 25 September, 2024. Seven (12%) problematic 2022 circRNA papers had been cited more than 50 times, with one paper (PMID 35768398) cited 106 times (Table S9). Problematic and other papers received comparable citation numbers, with a median of 21 citations per problematic paper (range: 4-106, n=60), versus 18 citations per paper for all other 2022 circRNA papers (range: 1-331, n=58) (Mann-Whitney Test, U=1,723, p=0.9284, n=118 papers).

### Post-publication commentary and corrections of problematic 2022 circRNA papers

Almost all problematic 2022 circRNA papers were indexed by PubPeer (58/60, 97%). As of 09 February 2026, a minority (14/58, 24%) had received PubPeer comments, which mostly (9/14) raised image integrity concerns. Two (3%) papers (PMID 35061934, 35210429) had PubPeer comments querying the identities of cell lines or nucleotide sequence reagents (Table S2), where the same errors were found in the present study.

During the study follow-up period (19 September 2023 - 09 February 2026), only 3/60 (5%) problematic 2022 circRNA papers (PMID 35361779, 35710754, 35061934) were retracted, and one paper (PMID 35210429) received a published correction (Table S2). All retraction notices described image integrity concerns that were previously raised on PubPeer (Table S2). One retraction referred to the use of HeLa contaminated cell lines, also identified in the present study (Table S2).

### Editorial handling and peer review of problematic 2022 circRNA papers

Reagent errors in publications could potentially reflect more rapid editorial and peer review processes. However, we found no significant difference in the median time to acceptance for problematic 2022 circRNA papers (median: 126.5 days, range: 16-405, n=60 papers) versus all other 2022 circRNA papers (median: 109 days, range: 10-298, n=58 papers) (Mann-Whitney test, U=1,409, p=0.0749, n=118).

## Discussion

Numbers of circRNA publications have increased markedly over the over the past decade (Wu et al. 2021; Sasso et al. 2022; Tuerdi et al. 2024; Yehni et al. 2024) (Figure 1). As we had previously found human circRNA papers describing with wrongly identified and NV nucleotide sequences in two high IF journals (Pathmendra et al. 2024), this study sought to verify reagent identities in circRNA papers across a broader range of similar journals.

We found wrongly identified reagents in all 13 retracted circRNA papers in high IF journals published between 2017-2021, where only one retraction notice described incorrect reagents. We subsequently found wrongly identified, NV and/or questionable reagents in 71% (84/118) 2022 circRNA papers in high IF journals, where 51% (60/118) papers described at least one wrongly identified reagent (Figure 3). Wrongly identified nucleotide sequence reagents were most frequent (44%), followed by questionable circRNA targeting probes (29%), NV nucleotide sequences (21%), wrongly identified cell lines (19%) and NV cell line identifiers (5%). Almost half of these papers described more than one type of reagent error or feature of concern.

### Study limitations

Before discussing these results, we should recognize that this study carries several limitations. We checked reagent identities in a minority of human circRNA papers, as we only examined papers in high IF journals. We recognise that journal IF reflects article citations, is easily manipulated (Siler and Larrivière, 2021) and does not reflect publication quality. Every paper was analyzed manually, which could produce false-positive results, particularly as thousands of individual reagents were analyzed. Some NV circRNA targeting reagents and questionable circRNA probes could reflect limitations of our fact-checking methods. Some NV circRNA reagents could reflect incomplete and non-overlapping circRNA databases and/or non-standardised circRNA nomenclature systems (Costa and Enguita, 2020; Dodbele et al. 2021; Vromman et al. 2021; Nielsen et al. 2022; Chen et al. 2023; Pathmendra et al. 2024). It may be difficult to predict whether some circRNA targeting reagents discriminate between linear and circular transcripts, based solely on the extents of sequence matches across BSJ sequences.

### Possible origins of reagent errors in circRNA papers

Reagent errors can arise in different contexts. Wrongly identified nucleotide sequence reagents can occur in genuine research (Park et al, 2022), particularly when papers describe many reagents. Nonetheless, some errors that this study described seem unlikely to be made by experts, such as claimed targeting reagents that do not appear to target any human gene or transcript. It is similarly difficult to explain circRNA targeting reagents that do not map to the relevant BSJ sequence, despite authors either providing the BSJ sequence or sufficient information to identify the claimed circRNA transcript. It was also surprising to find wrongly identified cell lines and NV cell line identifiers in articles in high IF journals, particularly as cell line descriptions would have been openly available (Capes-Davis et al. 2010; Bairoch, 2018) when these studies were conducted.

Many circRNA papers in this study were affected by multiple error types. For example, all 13 papers in the retracted circRNA corpus featured errors found by other investigators, typically image integrity errors described on PubPeer. We noted retracted papers in *Molecular Therapy-Nucleic Acids* and the *Journal of Experimental and Clinical Cancer Research* where undeclared involvement of a third party was described in retraction notices. Undeclared third party involvement could be synonymous with research paper mills (Han and Li, 2018), or unethical organisations that sell manuscripts and other publishing services (Byrne et al. 2022; Parker et al. 2024). *Molecular Therapy* journals have acknowledged being targeted by paper mills since 2020 (Bricker-Anthony and Giangrande, 2022; Frederickson and Herzog, 2022), also stating that circRNAs represent a preferred topic for paper mills (Bricker-Anthony and Herzog, 2023).

As journal IF partly dictates paper mill pricing (Abalkina, 2023), high IF journals could represent appealing targets for paper mills, where mistaken beliefs that high IF journals are not targeted by paper mills could increase their vulnerability (Pathmendra et al. 2024). Nonetheless, if paper mills are to successfully target high IF journals, they need to produce manuscripts that superficially resemble complex research studies. The regulatory functions of ncRNAs and circRNAs could provide layers of novelty and complexity to paper mill manuscripts that might meet the expectations of high IF journals (Pathmendra et al. 2024). Paper mills could also exploit non-standardised circRNA nomenclature and incomplete circRNA descriptions to render the detection of poor-quality circRNA research more challenging during editorial and peer review (Pathmendra et al. 2024).

### Implications of problematic circRNA papers in high IF journals

Most circRNA papers with wrongly identified reagents were published open-access, which would be predicted to be more widely read than articles behind paywalls (Huang et al. 2024; Yi et al. 2024). Problematic circRNA articles in many open-access high IF journals could encourage researchers to base new experiments on incorrect results. Researchers could also directly reuse incorrect or questionable reagents, leading to wasted resources and time. For example, a widely recognised circRNA function is “sponging” miRNAs, yet the combination of low circRNA abundance with few miRNA binding sites suggests that many circRNAs may not effectively sponge miRNAs or other regulators (Singh et al. 2024). As numerous retracted and problematic 2022 circRNA papers claimed to study circRNAs as “sponges” of miRNAs, the extent to which circRNAs function as miRNA “sponges” could be overestimated by unreliable publications.

### Next steps

Numbers of circRNA papers have risen more recently than papers focussing on other RNA topics such as miRNAs and lncRNAs (Sasso et al. 2022). This could explain why few studies have critically analysed reagent identities in circRNA publications (Patop and Kadener, 2018; Zhong et al. 2019; Oste et al. 2024; Pathmendra et al. 2024). Further analyses of circRNA papers in a broader range of journals and over different time periods could valuably inform the integrity of the human circRNA literature.

Prominent circRNA papers with wrongly identified, NV and questionable reagents support the need for community-led initiatives (Brumback et al. 2024) to promote integrity within the field. As noted in papers studying *miR-145* (Oste et al. 2024), the present study found that more circRNA papers described wrongly identified nucleotide sequences than cell lines, indicating that screening nucleotide sequence identities may be a sensitive means of identifying papers that could describe unreliable results (Byrne, 2025). Checking nucleotide sequence identities is taught to undergraduates (Niepielko and Shumskaya, 2021) and could be easily undertaken by most molecular researchers. The importance of also checking cell line identities was reinforced by almost all 2022 circRNA papers with wrongly identified and/or NV cell lines describing other errors and/or features of concern. Time investments in verifying reagent identities are therefore likely to be insignificant, compared with time potentially wasted on futile experiments.

This study also described low rates of post-publication corrections of 2022 circRNA papers. Only 3/60 problematic 2022 circRNA papers had been retracted and one paper had been corrected by February 2026, where only one retraction notice referred to any incorrect reagents. This low post-publication correction rate aligns with previous journal and publisher responses to nucleotide sequence errors (Grey et al. 2024). Journals and publishers need to act on the serious problems that wrongly identified reagents represent, as indeed recognised by Jung et al. (2000) more than 25 years ago. Prompt published flags of incorrect and NV reagents would help to protect readers from unreliable research (Byrne and Barnett, 2026). This is particularly important in the case of errors in open access papers in high IF journals that seem likely to attract attention, particularly from readers who mistake high journal IF for publication quality.

Our results also highlight opportunities for journals and publishers to impose higher standards on circRNA manuscripts and detect more verifiable errors prior to peer review. Many papers described circRNAs using nomenclature systems such as ArrayStar identifiers that do not allow circRNA identities to be verified. Mandatory circRNA descriptions that include BSJ sequences and genomic co-ordinates would permit independent verification of circRNA identities (Costa and Enguita 2020; Dodbele et al. 2021; Vromman et al. 2021; Nielsen et al. 2022; Chen et al. 2023; Pathmendra et al. 2024). Finally, descriptions of contaminated cell line models should be readily identifiable in submitted manuscripts, without access to specialist expertise (Byrne, 2025).

### Summary and conclusions

Our results show that many circRNA papers in high-IF journals describe the use of wrongly identified nucleotide sequences and/or cell lines. As such, we encourage researchers to exercise caution when reading the circRNA literature, including papers in high IF journals and with seemingly impressive citation numbers. Our results further support high IF journals as being potentially vulnerable to paper mill submissions. Given their recognition and available resources, we encourage high IF journals and their publishers to take the lead in responding to publications with wrongly identified and NV reagents (Pathmendra et al. 2024). Decisive actions from quality journals would set benchmarks for the broader community and play a key role in limiting the publication and impact of misleading circRNA research.

## Methods

### Identification of circRNA publication corpora

Retracted papers that referred to circRNAs in their titles were retrieved on 27 September 2023 using Web of Science as follows: keywords=“circular RNA*” (Title) or “circRNA*” (Title) or “hsa_circ*” (Title) or “hsa-circ*” (Title) or “hsacirc*” (Title) or “circ* RNA*” (Title), DT = “Retracted Publication”. We also included one retracted paper (PMID 32503552) that was found to describe wrongly identified reagents by Pathmendra et al. (2024).

Papers published in 2022 that referred to circRNA in their titles were retrieved on 19 September 2023 via Web of Science using the above keywords with additional modifications: PY = “2022”, DT = “Article”. Search results were exported to InCites (Clarivate), and journals and IFs were downloaded into Microsoft Excel. Journals with a 2022 IF≥7 were used as an additional filter for both retracted circRNA and 2022 circRNA corpora. Any 2022 circRNA papers that were retracted between 19 September 2023 and 09 February 2026 were included in the 2022 circRNA corpus, not the retracted circRNA corpus.

The titles of retracted circRNA and 2022 circRNA papers in high IF journals were used to query PubMed or the University of Sydney library to obtain publication PDFs and supplementary files. We also recorded the number of original circRNA papers published per year from 1976 to 2025, as reported on Web of Science on 15 April 2026.

### Visual inspection of circRNA papers

Articles were subjected to visual inspection and considered eligible for analysis if they described original experiments, studied at least one human circRNA, and either described the sequence of at least one nucleotide sequence reagent that was claimed to target a wild-type human transcript or genomic sequence, or the experimental use of at least one human cell line. Eligible papers were identified by PMID or otherwise by DOI.

### Verification of nucleotide sequence reagent identities

Nucleotide sequence reagents were checked according to their published status on 27 September 2023. Publication and supplementary files were inspected to determine the claimed identities of nucleotide sequence reagents and cell lines. Where 2022 circRNA papers had post-publication corrections, the identities of reagents in published corrections were also examined.

Claimed identities of targeting nucleotide sequence reagents were checked using information in GeneCards (Stelzer et al. 2016), miRBase (Griffiths-Jones et al. 2006), lncATLAS (Mas-Ponte et al. 2017), circBase (Glažar et al. 2014) and circAtlas (Wu et al. 2020). GeneCards (Stelzer et al. 2016), miRBase (Griffiths-Jones et al. 2006), and Ensembl (Harrison et al. 2024) were used to clarify synonymous identifiers and identify chromosomal positions of claimed identifiers. In the absence of any clearly linked reagent identifier, reagent identities could sometimes be determined by analysing the context in which reagents were used. If any claimed reagent identity was not evident or if reagents were claimed to target non-human genes or if cell lines represented non-human models, these reagents were excluded from analysis.

Nucleotide sequence reagents were analysed using Blastn (Altschul et al. 1990), BLAT against the GRCh38/hg38 assembly (Lee et al. 2022), or the alignment tool available on miRBase (Griffiths-Jones et al. 2006) as described (Park et al. 2022; Pathmendra et al. 2024). Reagent sequences claimed to target human miRs were also manually aligned with miR transcripts available on miRbase, as described (Park et al. 2022). Reagents claimed to target long non-coding RNAs (lncRNAs) were verified as described by Pathmendra et al. (2024). Where necessary, reagent sequences were reversed or reverse-complemented using the Sequence Manipulation Suite (Stodhard, 2000). Wrongly identified reagents were classified according to previously described error categories (Byrne et al. 2021; Park et al. 2022; Pathmendra et al. 2024).

### Verification of claimed circRNA targeting reagent identities

We first checked whether claimed circRNA transcripts could be identified through the disclosure of any circRNA identifier or information such as circRNA length, BSJ sequence and/or chromosomal location that could be queried using circPrimer (Zhong et al. 2018), circBase (Glažar et al. 2014) or circAtlas (Wu et al. 2020). If the claimed circRNA transcript could not be identified, associated reagents were described as non-verifiable (NV) (Pathmendra et al. 2024).

Claimed convergent RT-PCR primers were verified as RT-PCR primers for linear transcripts (Dudekula et al. 2016; Zhong et al. 2018; Nielsen et al. 2022; Pathmendra et al. 2024). Claimed divergent RT-PCR primers were initially queried on circPrimer (Zhong et al. 2018) as described (Pathmendra et al. 2024). If circPrimer produced no output, primer sequences were manually to the forward and reverse complement sequences of the claimed circRNA transcript on either side of the BSJ.

circRNA knockdown reagents typically target the circRNA BSJ (Dudekula et al. 2016; Nielsen et al. 2022) and were verified as described (Pathmendra et al. 2024). If a circRNA knockdown reagent showed 100% identity over ≥ 17 consecutive nucleotides with linear transcripts, including from the claimed host gene, the reagent was classified as wrongly identified, as such reagents may not discriminate between circular and linear isoforms. Wrongly identified circRNA targeting reagents were classified as described by Pathmendra et al. (2024).

### Verification of circRNA reagent identities used in F/ISH, RNA pull-down and Northern blot analyses

circRNA targeting reagents used in F/ISH, RNA pull-down or Northern blot experiments were assumed to target the BSJ of the claimed circRNA, unless stated otherwise. We verified whether reagents satisfied previously described targeting criteria (Table S6) and recorded the lengths of sequence matches on each side of the BSJ in MS Excel. Reagents that were claimed to target the BSJ but predicted to match sequences on only the 5’ or 3’ side of the BSJ were noted as wrongly identified.

Reagents that matched both sides of the claimed BSJ but did not meet specific targeting length requirements (Table S6) were described as questionable.

### Verification of human cell line identities

Human cell line identities were verified as described by Oste et al. (2024). Briefly, if the claimed cell line identity did not match the verified identity on Cellosaurus (Bairoch, 2018), the cell line was classified as wrongly identified. Human cell lines previously described to be cross-contaminated by a different cell line or misclassified were flagged as problematic, if papers did not acknowledge i) the cell line’s contaminated/ misclassified status or ii) use of a contaminated/ misclassified cell line as a study limitation.

If any cell line identifier was not indexed by Cellosaurus, the identifier was queried on the American Type Culture Collection catalogue or any externally accessible source catalogue described by the study. If these approaches produced no results, the identifier was queried on Google Scholar combined with terms such as “cells” or “cell line” to identify the earliest publication(s) that would be expected to describe cell line establishment. If no establishment publication could be found, further keyword searches were conducted on Cellosaurus and Google Scholar, with variations in the spelling of the identifier to identify a possible misspelled origin of the NV cell line (Oste et al. 2024). If no evidence of cell line establishment could be found through these combined approached, the cell line identifier was described as non-verifiable (NV).

### Publication classification

circRNA papers were classified as problematic if they included at least one wrongly identified nucleotide sequence reagent and/or one wrongly identified cell line. Papers that only described NV reagents and/or questionable circRNA targeting reagents were recorded but not classified as problematic.

### Additional publication analyses

Publication titles were visually inspected for mentions of specific circRNAs, human cancer types, the terms “sponging” and/or “axis”, and drug identifiers, which were confirmed through Google searches. Countries of origin and institutional affiliations were classified as described (Park et al. 2022). Where there was no numeric majority, the first author’s affiliation was used to decide the overall institutional affiliation and/or country of origin (Pathmendra et al. 2024). Citation numbers according to Google Scholar were obtained on 25 September, 2024. Dates of manuscript submission and acceptance were obtained from publication webpages and the difference (in days) was calculated as time to manuscript acceptance. The open access status of 2022 circRNA papers was obtained by visually inspecting publication webpages.

### Analyses of retraction notices and PubPeer commentary

Retraction notices were visually inspected for information about which party/ies had initiated and/or authorised retractions and coded based on the reason(s) for retraction. Times to retraction were calculated by subtracting publication dates from retraction dates (in months/ years). Proportions of retracted circRNA papers were noted according to (i) reason(s) for retraction, (ii) whether the stated reason(s) for retraction aligned with any PubPeer comments made before the retraction date and (iii) party(ies) who initiated the retraction.

Where circRNA papers were retracted due to concerns described in earlier PubPeer posts, times to retraction from the first PubPeer query were calculated by subtracting the date of the earliest PubPeer post that was addressed in the retraction notice from the retraction date (in months). PubPeer comments for problematic 2022 circRNA papers were visually inspected to summarise the number and content of PubPeer posts, whether publication authors had responded to PubPeer comments, and dates of PubPeer posts (month/ year).

### Statistics

Spearman’s rank correlation compared numbers of wrongly identified nucleotide sequences with numbers of verified nucleotide sequences per problematic 2022 circRNA paper. The Mann-Whitney test was used to compare article citations and times to acceptance, according to 2022 circRNA publication error status. Fisher’s Exact test compared the proportions of 2022 circRNA papers according to error status, and country or institutions of origin. Statistical analyses were performed on GraphPad Prism 10. Graphs were produced on MS Excel or GraphPad Prism 10. Reported p values have not been corrected for multiple comparisons.

## Acknowledgements

We thank Dr Amaresh C. Panda (Institute of Life Sciences, Bhubaneswar, India) and Prof. Argyris Papantonis (University Medical Center of the University of Göttingen, Germany) for valuable insights and discussions about the targeting parameters of F/ISH and RNA pull-down reagents.

